# Transcriptional signatures of clonally derived *Toxoplasma* tachyzoites reveal novel insights into the expression of a family of surface proteins

**DOI:** 10.1101/2021.12.26.474199

**Authors:** Terence C. Theisen, John C. Boothroyd

## Abstract

*Toxoplasma gondii* has numerous, large, paralogous gene families that are likely critical for supporting its unparalleled host range: nearly any nucleated cell in almost any warm-blooded animal. The *SRS* (SAG1-related sequence) gene family encodes over 100 proteins, the most abundant of which are thought to be involved in parasite attachment and, based on their stage-specific expression, evading the host immune response. For most SRS proteins, however, little is understood about their function and expression profile. Single-parasite RNA-sequencing previously demonstrated that across an entire population of lab-grown tachyzoites, transcripts for over 70 *SRS* genes were detected in at least one parasite. In any one parasite, however, transcripts for an average of only 7 *SRS* genes were detected, two of which, *SAG1* and *SAG2A*, were extremely abundant and detected in virtually all. These data do not address whether this pattern of sporadic *SRS* gene expression is consistently inherited among the progeny of a given parasite or arises independently of lineage. We hypothesized that if *SRS* expression signatures are stably inherited by progeny, subclones isolated from a cloned parent would be more alike in their expression signatures than they are to the offspring of another clone. In this report, we compare transcriptomes of clonally derived parasites to determine the degree to which expression of the SRS family is stably inherited in individual parasites. Our data indicate that in RH tachyzoites, *SRS* genes are variably expressed even between parasite samples subcloned from the same parent within approximately 10 parasite divisions (72 hours). This suggests that the pattern of sporadically expressed *SRS* genes is highly variable and not driven by inheritance mechanisms, at least under our conditions.

## Introduction

*Toxoplasma* is a ubiquitous, intracellular, eukaryotic pathogen that can cause severe illness in immunocompromised individuals and the developing fetus. *Toxoplasma*’s lifecycle includes sexual stages that only arise and mate in the intestines of felines, and asexual stages that infect and reproduce in virtually any nucleated cell across a very large number of warm-blooded animals (1, 2). The asexual stages include rapidly growing tachyzoites and nearly quiescent bradyzoites, which form persistent cysts in muscle and brain tissues of an infected host. *Toxoplasma* can transmit to new hosts through the ingestion of cyst-containing tissue (3, 4). The unparalleled host range of *Toxoplasma* is thought to increase the likelihood of exposure to a cat intestine due to the indiscriminate carnivorous habits of felines (5, 6). The molecular mechanisms that support *Toxoplasma*’s ability to attach to, infect, and successfully evade the immune responses of many different cell types within numerous mammalian and avian species are only just beginning to be understood.

Members of a large family of glycosylphosphatidylinositol (GPI) anchored surface antigens, the SRS (SAG1-related sequence) family, have been proposed to be involved in *Toxoplasma*’s attachment to host cells and immune evasion (7, 8). For example, evidence exists to indicate SAG1 (Surface Antigen 1, also known as SRS29B; note that throughout this manuscript, where more than one name exists we will use the historic gene/protein names for simplicity while providing both names on first use) and SAG3 (SRS57) play a role in attachment, while SAG1 and SAG2A (SRS34A) have been suggested to influence the host immune response (9–13). SRS2 (SRS29C) has been shown to vary in its expression levels between strains and to contribute to overall virulence of the parasite in a mouse model (8). All *SRS* genes encode at least one SRS-fold, a protein domain of about 20 kDa that forms a unique structure through disulfide bonds between 4 or 6 conserved cysteine residues and is only observed in Apicomplexan species (14). Crystal structures and modeling of SRS members revealed the likely formation of homodimers with a groove of positively charged residues that has been proposed as a binding region for negatively charged polymers, possibly glycosaminoglycans (GAGs) on a host cell (14). Interestingly, across the *SRS* gene family, there is greater variation in the protein sequence of the proposed binding domains compared to the more conserved regions that are proximal to the C-terminal GPI-anchor (8).

Early protein-based studies demonstrated that some SRS family members are expressed in a stage-specific manner and recent RNA sequencing studies on the various life stages of *Toxoplasma* have further supported this finding (15–19, 21). There are ~10 SRS proteins that are abundant in tachyzoites (e. g., SAG1, SAG2A, SRS2, SRS3 (SRS51), SAG3, SRS20A, SRS25, SRS1 (SRS29A), SRS52A, SRS67) while a nearly non-overlapping set is abundantly present in bradyzoites (e.g., SRS44 (CST1), SRS9 (SRS16B), SAG2X (SRS49B), SAG2Y (SRS49A), SAG2C (SRS49D)), sporozoites (SporoSAG (SRS28)), and the sexual stages found within the cat intestine (SAG2B (SRS11), SRS22A/B/C/D/E/F/G/I, SRS30A/C/D, SRS48E/K/Q) (16, 17, 19, 22–24). In addition to these highly expressed members, transcriptomic and proteomic data from analysis of bulk populations showed many additional SRS family members are expressed at very low levels in these developmental forms (15–21, 25). With all these data, however, it was unclear whether these low-abundance SRSs are consistently expressed at low levels across the entire population of parasites or if a minority of parasites express different subsets of *SRS* genes at somewhat higher levels while a majority do not express them at all.

Recent work from our lab and others characterizing the single parasite transcriptomes of the asexual stages of *Toxoplasma* shows that in any given tachyzoite of the RH strain, transcripts for an average of just 7 *SRS* genes are detected, whereas across the entire tachyzoite population, transcripts for nearly 70% of *SRS* genes are detected in at least one parasite (26, 27). As expected, transcripts for the highly abundant *SAG1* and *SAG2A* are detected in more than 99% of parasites, while of the remaining 83 detected SRS family members, 11 are detected in between 10 and 99% of parasites, 29 are detected in 5-10% of parasites and 43 are detected in less than 5% of parasites. Only a small number (<8) of *SRS* genes appear to be cell-cycle regulated, and so this could not account for the variation seen in *SRS* transcript expression (18, 26). Likewise, controls using spiked in, synthetic RNAs of known relative abundance showed that the failure to detect most SRS genes was not a technical artifact due to low sensitivity (26). These data strongly suggested, therefore, that in a non-clonal population of tachyzoites there is a significant degree of variation in the expression of low-abundance *SRS* genes, rather than a generally low level of expression across the entire population. These data did not, however, address whether this pattern of “sporadic” *SRS* gene expression is consistently inherited among the progeny of a given parasite, or whether expression of these genes is stochastic, resulting in a pattern that is independent of lineage. To address this, we report here transcriptomic data on tachyzoite samples approximately 10 generations after they have been isolated from a common parent, thereby providing insight into the degree to which a given pattern of sporadic *SRS* transcript expression is inherited. Our results provide evidence that the expression pattern of sporadically expressed *SRS* genes is not stably inherited among clonally related parasites, supporting the notion that their expression in tachyzoites, at least, is stochastic.

## Materials and Methods

### Parasite preparation

A population of *Toxoplasma gondii* RH expressing mCherry (26) was used for all experiments described here. The precise number of passagings every ~3 days since this line was generated and last cloned is not known but it is at least 90, thereby representing over 600 generations (doublings) of the parasite.

### Limiting dilution to isolate clones and subclones

Human foreskin fibroblasts (HFFs) were seeded in 96 wells plates and grown to confluency in DMEM medium (ThermoFisher 11960044) at 37°C in 5% CO (26). RH mCherry^+^ parasites were scraped and syringed-released from an overnight culture of a heavily infected T25 flask of HFFs, passed through a 5 μm filter (ThermoFisher/Millipore SLSV025LS), and counted. For the first limiting dilution, parasites were added to multiple plates at concentrations yielding 1 parasite/well or 2 parasites/well. Plates were spun down at 800 rpm for 5 minutes at 4°C and placed at 37°C. To limit the potential for multiple infections in one well, after 2 hours the infection media was removed, new media was added, plates were spun again as above, and returned to the 37°C incubator. One plate was set aside for clone harvesting (see below).

For subclone isolation a second limiting dilution was performed thirty hours after the first limiting dilution and growth at 37°C. Wells containing clones were examined visually with brightfield light and illumination to detect mCherry at 20x magnification to identify wells that contained only one parasite vacuole with a size of 8, 16, or 32 parasites. Single-vacuole wells were scraped thoroughly with a pipet tip and then the entire volume of ~200 μl passed through a 27 gauge needle (VWR B27-50) at least 4 times. The volume of the syringe-released solution was expanded and added to 24 wells of a new 96-well plate of confluent HFFs. The plate was spun down at 800 rpm for 5 minutes at 4°C and then incubated at 37°C.

### Clone and subclone sample harvesting

Clones were harvested seventy-two hours after the first limiting dilution and subclones were harvested seventy-two hours after the second limiting dilution. As for generation of the clones, the full area of a well containing putative subclones was visually inspected with both brightfield and UV fluorescence at 20x magnification to identify wells with only one apparent focus of infection. To collect parasites, a pipet tip was used to disrupt and loosen the HFF monolayer. A well’s content was passed at least 4 times through a 27 gauge needle. Due to the low volume of the well contents (< 200 μL) a 5 μm filter was primed with 400 μL of PBS. As the well contents flowed by gravity through the filter, an additional 200 μL of PBS was added to the syringe to flush the filter. The solution was collected in a 1.5 mL tube. The samples were then spun at 15,000 g for 10 minutes at 4°C and the supernatant gently removed by pipet. Samples were then resuspended in 10 μL of a solution with 200 U of RNase I (ThermoFisher/Ambion AM2295) for 20 minutes at 37°C. Directly after the incubation, 100 uL of TRIzol (ThermoFisher 15596026) was added to the samples. Samples were kept at −80°C until further processing.

### RNA extraction and cDNA synthesis

Samples in TRIzol were thawed on ice and all RNA was extracted using a standard chloroform/isopropanol RNA purification, per the manufacturer’s instructions. Following the final ethanol wash, pellets were resuspended in 4 μL of Pre-RT buffer (RNase-free water (ThermoFisher 10977023), dNTPs (ThermoFisher R0194; 2.5 mM), and RNA inhibitor (TaKaRa 2313; 10 units/reaction), Oligo-dT (IDT; 2.5 μM)) except for sample C3c, which was resuspended in 8 uL of Pre-RT buffer and divided into two aliquots. For reverse transcription, 6 μL of Reverse Transcriptase Buffer (5x First strand buffer (TaKaRa 639538), Betaine (Sigma B0300; 1 M), DTT - Dithiothreitol (TaKaRa 639538; 1.66 mM), TSO - Template Switch Olgio (IDT; 1 μM), MgCl_2_ (ThermoFisher AM9530G; 7 mM), and SmartScribe Reverse Transcriptase (TaKaRa 639538; 100 units/reaction)) were added to each of the samples (28). Per the instructions for the reverse transcriptase kit, the samples were run on a thermocycler with the following steps: 1) 42°C for 90 minutes; 2) 50°C for 2 minutes; 3) 42°C for 2 minutes; 4) repeat steps 2 and 3 (10x); 5) 70°C for 15 minutes. Next, cDNA was amplified using HiFi Hotstart Ready Mix (ThermoFisher 50-196-5217) with ISPCR (IDT; 0.1 μM) and Lambda Exo (New England BioLabs MO262S; 1 units/reaction) for 12 or 14 total cycles.

### Library Preparation, pooling, and sequencing

Amplified cDNA was purified using AMPure XP beads (Beckman Coulter A63881) and concentration was determined by Qubit 4.0 using the 1x dsDNA HS kit (ThermoFisher Q33230). Nextera XT indices were used to generate libraries from 1 ng of cDNA material for each sample following the instructions provided by Illumina (Illumina FC-131-1096 and FC-131-2001). Library preparations were purified using AMPure XP beads and checked for quality using the Agilent Bioanalyzer at Stanford Functional Genomics Facility (SFGF). A sequencing pilot was done with libraries pooled in equal amount and run on one NextSeq500 lane with high output, 1×75 sequencing. Using the results from this sequencing a second library was prepared based on the fraction of unique *Toxoplasma* reads to total reads for a library in an attempt to generate approximately the same number of *Toxoplasma* reads per sample. The second library was run over two lanes NextSeq500 lanes with high output, 1×75 sequencing at the SFGF. Results from the different lanes for each clone or subclone were pooled prior to alignment.

### Sequencing alignment

Read outputs from sequencing were aligned using STAR aligner (Version 2.7) to a concatenated human (GRCh38.p13 accessed March 2021) and Toxoplasma ME49 genome (ToxoDB v51 accessed March 2021) (15, 29). Transcript counting was performed using HTSeq-count with the same parameters used in Xue (26, 30).

### Expression analysis

*Toxoplasma*-specific read counts were filtered from the HTSeq-count dataset. Read counts were normalized using the median read sum of the samples’ uniquely counted *Toxoplasma*-specific reads. Down-sampling was done using pandas sample method for 5 randomly chosen random states (3, 6, 10, 185, and 278). The clustermap was generated using seaborn’s clustermap function using ‘average’ as the linkage method and ‘euclidean’ as the distance metric.

### Statistical analysis

Data processing and analysis was done in Python 3.8.8. Packages used in the analysis included matplotlib (3.3.5) (31), matploblib-venn (0.11.6), numpy (1.20.1) (32), pandas (1.2.4) (33), scikit-learn (0.24.1), and seaborn (0.11.1) (34). R (4.1) (35) and the DESeq2 (1.32.0) (36) library were used for the differential expression analysis.

## Results

### Experimental approach

To study whether *SRS* gene expression signatures are inherited, we performed RNA sequencing on tachyzoites that had recently been cloned (i.e., expanded from a single parasite obtained through limiting dilution; Fig 1). These parasites were derived from a starting population of mCherry^+^ RH parasites that had not been cloned in at least 90 passages (representing at least 600 divisions based on passage every 2 to 3 days in human foreskin fibroblasts (HFFs)). This passage history is similar to that of the tachyzoites used in the single-parasite RNA-sequencing study (26) where we saw extensive differences in the repertoire of *SRS* expression in individual cells. Individual parasites within this lab-passaged population were cloned by limiting dilution into 96-well plates harboring HFFs. Thirty hours after this cloning, “subclones” of selected clones were isolated through a second limiting dilution of the parasites present in a single well. Only clones (C1, C2, etc.) and subclones (C1a, C1b, C1c, C2a, C2b, etc.) that were visually confirmed to be derived from a single parasite, taking advantage of the fluorescent mCherry expressed by these parasites, were used in subsequent analyses.

**Fig 1.**
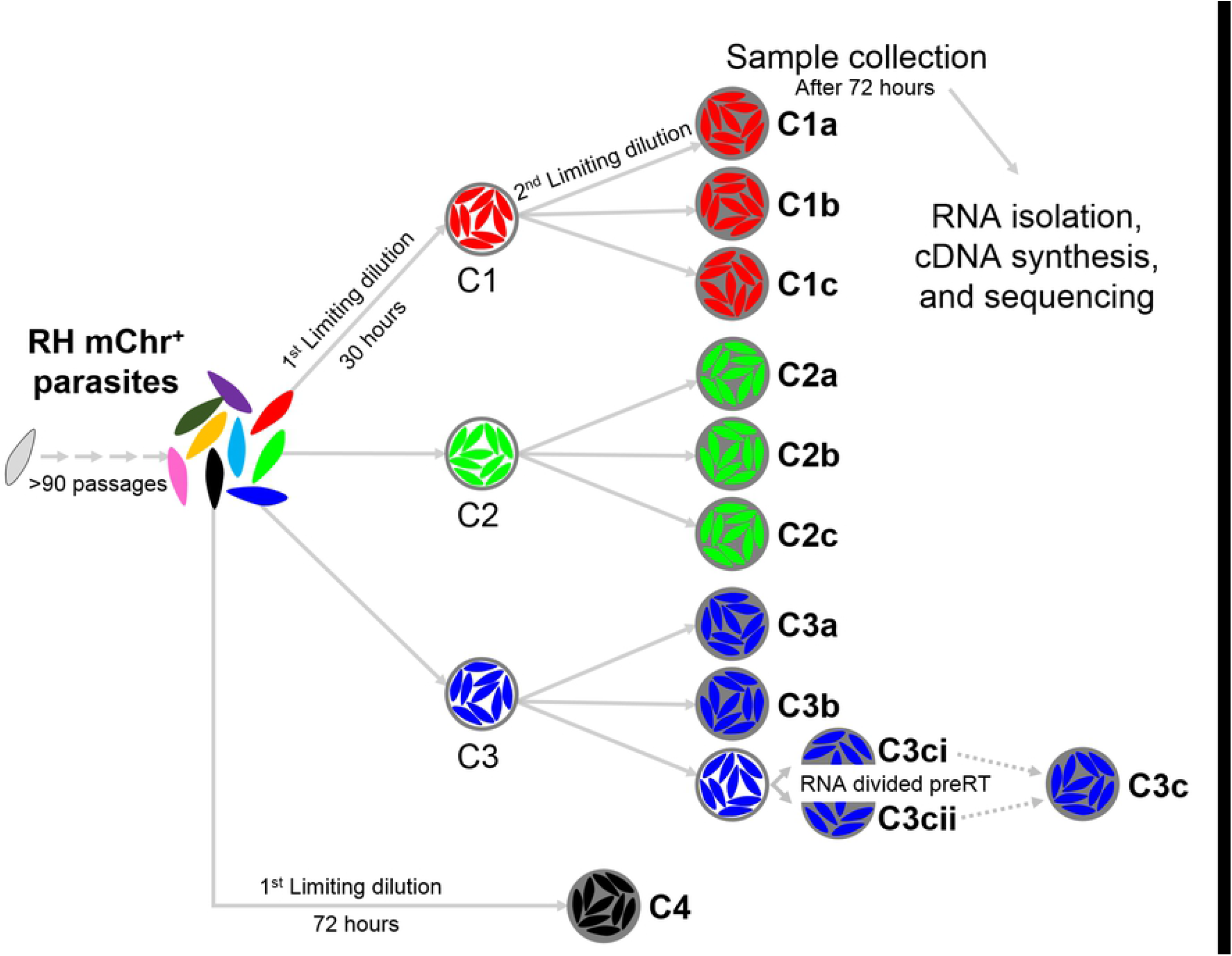
*Toxoplasma* subcloning schematic. A population of mCherry^+^ (mChr^+^) RH parasites was passed at least 90 times in human foreskin fibroblasts (HFFs) after thawing. Independent clones (indicated by colors and given the names C1, C2, C3, and C4) from the population were isolated by a first limiting dilution into 96-well plates with an HFF monolayer. After thirty hours, three wells with a single vacuole were identified and individual parasites were isolated by a second limiting dilution into new 96-well plates. Seventy-two hours after the second limiting dilution, subclones samples (designed by the lowercase letters) were harvested from wells for RNA sequencing (filled circles). A clone, C4, from the original population was also collected seventy-two hours after the first limiting dilution to mimic the collection of the subclones (same time from isolation of a single parasite in a well to harvest). For subclone C3c, the extracted RNA was divided prior to the reverse transcriptase reaction to yield C3ci and C3cii, and in some analyses these two datasets were aggregated to yield the “C3c” sample.

Harvesting of subclones occurred ~72 hours after the second limiting dilution. In preliminary experiments we determined that this growth period yields ~500-2000 parasites, about the number expected for 3 days of outgrowth from a single parasite and representing ~10 divisions. Additionally, while the single-parasite transcriptomic data did not indicate that *SRS* genes, as a group, were strongly correlated to a parasite’s cell cycle state, allowing ~10 divisions mitigated any such effect by allowing enough expansion that the parasites were dividing asynchronously (such synchrony is lost after one lytic cycle or about 48 hours) (18). On the other hand, we wanted to limit the number of generations that occurred after subcloning to increase our chances of seeing stable inheritance of *SRS* gene expression, should such be occurring. Because we are interested here only in the parasite’s transcriptome, we syringe-lysed the infected cells in wells containing a single clone or subclone, to release intact parasites and then passed this material through a 5 µm filter which allows tachyzoites to pass while removing unlysed host cells and much host cell debris. Gel electrophoresis with an agarose gel of the total RNA showed that this technique results in a substantial reduction in total host cell RNA based on loss of the host 28S large rRNA (Fig 2). Following filtering, the parasites were spun down and treated with RNase I with the aim of further reducing host cell RNA before resuspension in TRIzol.

**Fig 2.**
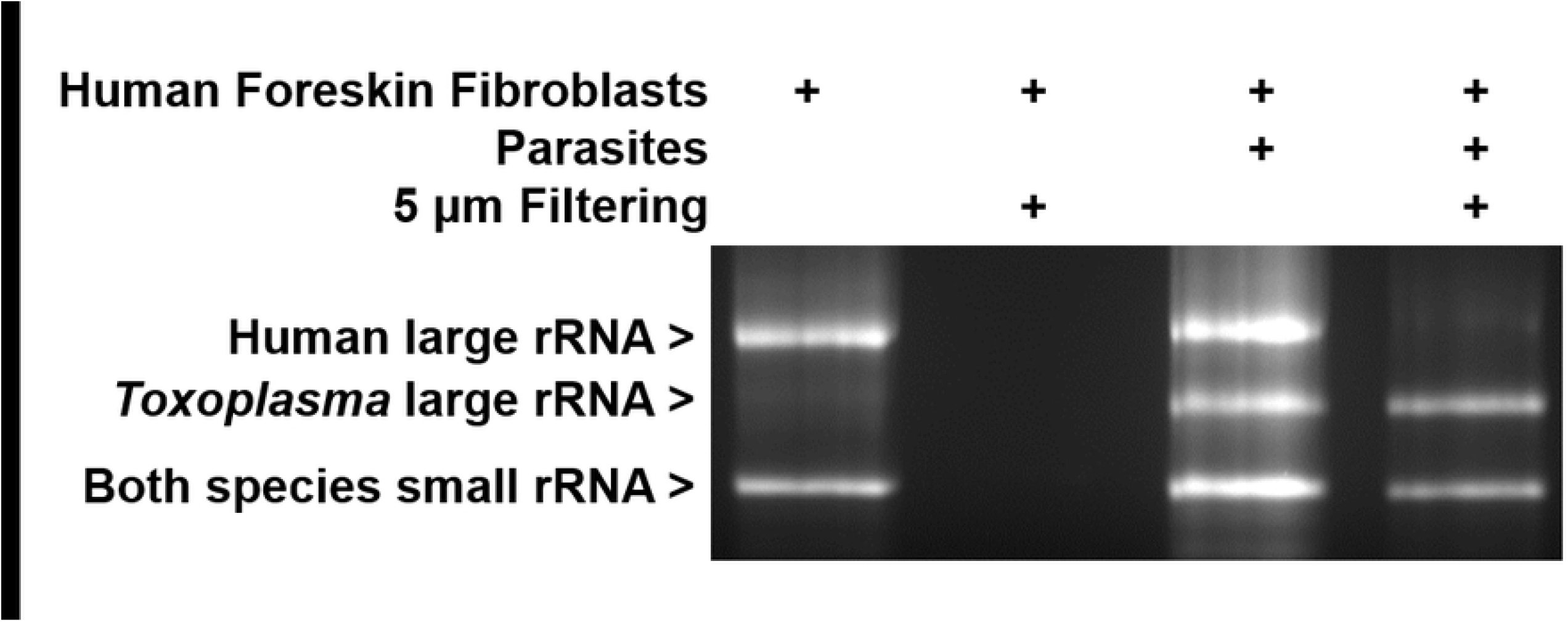
Filtering reduces host RNA. T25 flasks with HFFs (human foreskin fibroblasts) were infected with tachyzoites or left uninfected. Contents of a flask were scraped, passed through a 27 g needle three times, and half was then passed through a 5 μm filter to remove host material, the other half being kept for comparison. RNA was extracted from all samples using TRIzol and analyzed by agarose gel electrophoresis with visualization using ethidium bromide staining. Bands corresponding to the human and *Toxoplasma* large and small subunit rRNAs are indicated.

Prior to reverse transcription (RT), the RNA for one of the C3 subclone samples was divided into two aliquots to serve as a technical replicate control (C3ci and C3cii). All samples were prepared for RNA sequencing using the Nextera XT Library preparation kit and the libraries were run on the NextSeq500 platform following a protocol adapted from Rastogi et at (37). Reads were aligned to concatenated human and *Toxoplasma* ME49 genomes using the method described in Xue et at (26). Across the samples, reads that mapped uniquely to *Toxoplasma* transcripts accounted for between 0.5% and 3.7% of the reads that uniquely mapped to human or *Toxoplasma* transcripts. In the wells that subclones were collected from, HFF cells outnumbered tachyzoites by 30 to 60-fold. Given that there are approximately 500,000 to 1,000,000 mRNA molecules per human cell vs. approximately 40,000 to 50,000 for *Toxoplasma* tachyzoites (26, 38), we estimate that the parasite mRNA in these samples started out being <0.03% of the total mRNAs present. This indicates that the filtering and/or RNase treatments resulted in at least a 15-fold enrichment for *Toxoplasma* transcripts.

### *Toxoplasma* gene expression in clonal samples

To determine if the low percentage of *Toxoplasma* transcripts relative to total transcripts resulted in limited detection of *SRS* genes, we started by examining the number of *Toxoplasma* genes detected across our samples. The ME49 genome was used as the reference genome due to the higher number of annotated genes (39). The total uniquely aligned *Toxoplasma* read sums across the samples ranged from 61,544 to 371,843 (S1 Table). Given this range, we wanted to determine if our samples had comparable gene detection levels. Samples were normalized by dividing each read count by the read sum of the corresponding sample and then multiplying by the median read sum of the samples in the dataset to yield count per median (CPM) as in Xue et al (26) (S2 Table). We then calculated the number of genes detected above 4 CPM for each sample and plotted against the corresponding read sum (Fig 3 top). This value of 4 CPM was chosen to ensure that a gene was not incorrectly considered to be expressed due to an aberrant sequence read. Across the range of *Toxoplasma* read sums the number of genes detected above 4 CPM was generally between 2900 and 3700, regardless of the total number of reads. The linear regression value (r^2^) of the read sums and genes detected above 4 CPM was 0.035, indicating essentially no correlation between the two measures. This suggests the number of detectable genes is saturated at the level of sequencing coverage in the experiment.

**Fig 3.**
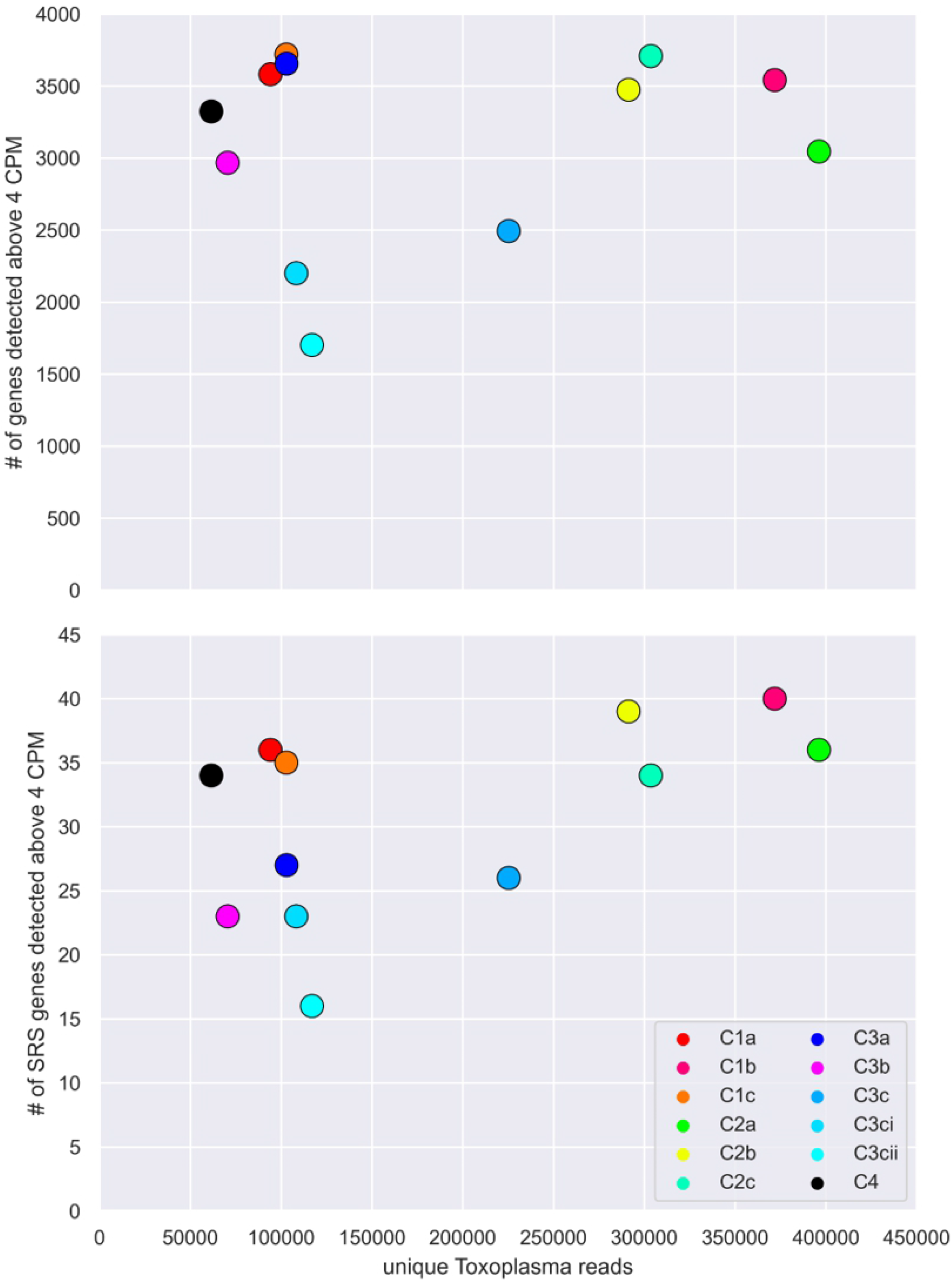
*Toxoplasma* transcript detection and read depth. For each sample, the total number of *Toxoplasma* genes “detected” in that sample is plotted against the total number of reads that uniquely aligned to the *Toxoplasma* genome in that sample (top panel). A gene was considered “detected” if it was above 4 counts per median (CPM; see Materials and Methods). C3ci and C3cii represent the results from the RNA sample that was split prior to cDNA synthesis. C3c represents the aggregation of the C3ci and C3cii data. Bottom panel is the same analysis as the top panel except only reads mapping to the 111 annotated *SRS* genes were counted for plotting on the y-axis.

Two of the three exceptions to the lack of correlation between a sample’s read sum and genes detected were the two aliquoted samples, C3ci and C3cii, from the C3 subclone which was split prior to the reverse transcription step. When bioinformatically combined to mimic the result we would have presumably obtained if the C3c sample had not been split, 2493 genes were detected. This compares to a range of 2900-3700 detected genes for the unsplit samples. For C3ci and C3cii, however, only 2200 and 1702 genes were detected, respectively (Fig 3 top), with 1077 genes detected in both samples, 1123 genes detected in only C3ci relative to C3cii, and 625 genes detected in only C3cii relative to C3ci. Because these were aliquots of the same RNA preparation and all of the 1748 genes whose transcripts were detected in only one or other aliquot were detected in at least one other sample, we conclude that, not surprisingly, given the very low amounts of starting material and over-abundance of host material, the protocol we are using fails to efficiently capture very low abundance parasite mRNAs. The way the C3c replicates were processed may have contributed to this observation: the RNA from the C3c well was resuspended in twice the volume of reverse transcriptase buffer as the other samples and split into two separate wells (C3ci and C3cii) for the steps after this point. The data would suggest that increased dilution of the *Toxoplasma* RNA from the well resulted in lower overall capture efficiency of the possible mRNAs. The number of *Toxoplasma* genes detected in the other samples is similar to the number of genes detected by RNA sequencing in a population of asynchronous tachyzoites, previously reported to be ~2000-4000 bulk studies (16, 17).

### *SRS* expression signatures vary between subclones

When considering just the 111 annotated *SRS* genes in the ME49 genome, 65 were detected above 4 CPM in at least one of the clones. In the C1- and C2-originating subclones, between 33 and 40 *SRS* genes were detected (Fig 3 bottom). The C3 subclones had substantially lower counts of detected *SRS* genes ranging from 23 to 27 (using the pooled “C3c” data for C3ci and C3cii). The r^2^ value of the linear regression using the SRS detection vs. total *Toxoplasma* reads (Fig 3) is higher (0.288) than the r^2^ for the same analysis using the total genes detected (0.05; Fig 3). The lower number of *SRS* genes detected in the C3a and C3b subclones is not a reflection of a lower number for the detection of all *Toxoplasma* genes, indicating this result is specific to this gene family and suggesting that there is some level of control being exerted on expression of at least a subset of *SRS* genes. Given that many of the sporadically expressed *SRS* genes detected in Fig 3 are most abundantly expressed in stages other than tachyzoites (17, 24), these data may indicate that the C3 subclones are more tightly locked into the tachyzoite state than the other cloned lines. Consistent with this hypothesis is the fact that, for example, all three C1 subclones express substantial levels of two canonical bradyzoite *SRS* genes, SRS9 (40) and CST1 (41), while their expression in the C3 subclones is below the threshold of 4 CPM. This point is discussed further below.

We hypothesized that if *SRS* expression signatures are stably inherited by progeny, subclones from the same parent clone would be more similar to each other, with regard to their expression signatures, than they are to the offspring of another clone. To test this, we first determined whether any *SRS* genes were differentially expressed between the clonal lineages by considering each subclone sample as a biological replicate of a clonal lineage. We used DESeq2 to compare differentially expressed genes between the clonal lineages (S3 Table) (36). Only twenty genes showed evidence of differential expression (adjusted p-value < 0.05), based on the pairwise comparisons between the different clones: C1 vs C2, C1 vs C3, and C2 vs C3 (Fig 4), and of these 20, none were *SRS* genes and only 7 had an adjusted p-value < 0.01. We also did the differential expression analysis where the replicates were grouped not on lineage but on a random variable (i.e., comparing all “a” vs. all “b”, all “a” vs. all “c”, and all “b” vs. all “c” replicates); by this comparison only two genes, 267460 (AP2IXI) and 314750, had adjusted p-values of less than 0.05. The differential expression of AP2IX1 is entirely due to its detection in just one sample, C1a, with 48 counts compared to 0 counts in all other samples. This argues that differences in expression of these 20 genes across the three parasite clonal linages might be real differences, though whether they are stable beyond the duration tested here would require further testing. The absence of a known function for all but one gene in this list makes it difficult to comment on the biological impact of, or possible reason for, these differences.

**Fig 4.**
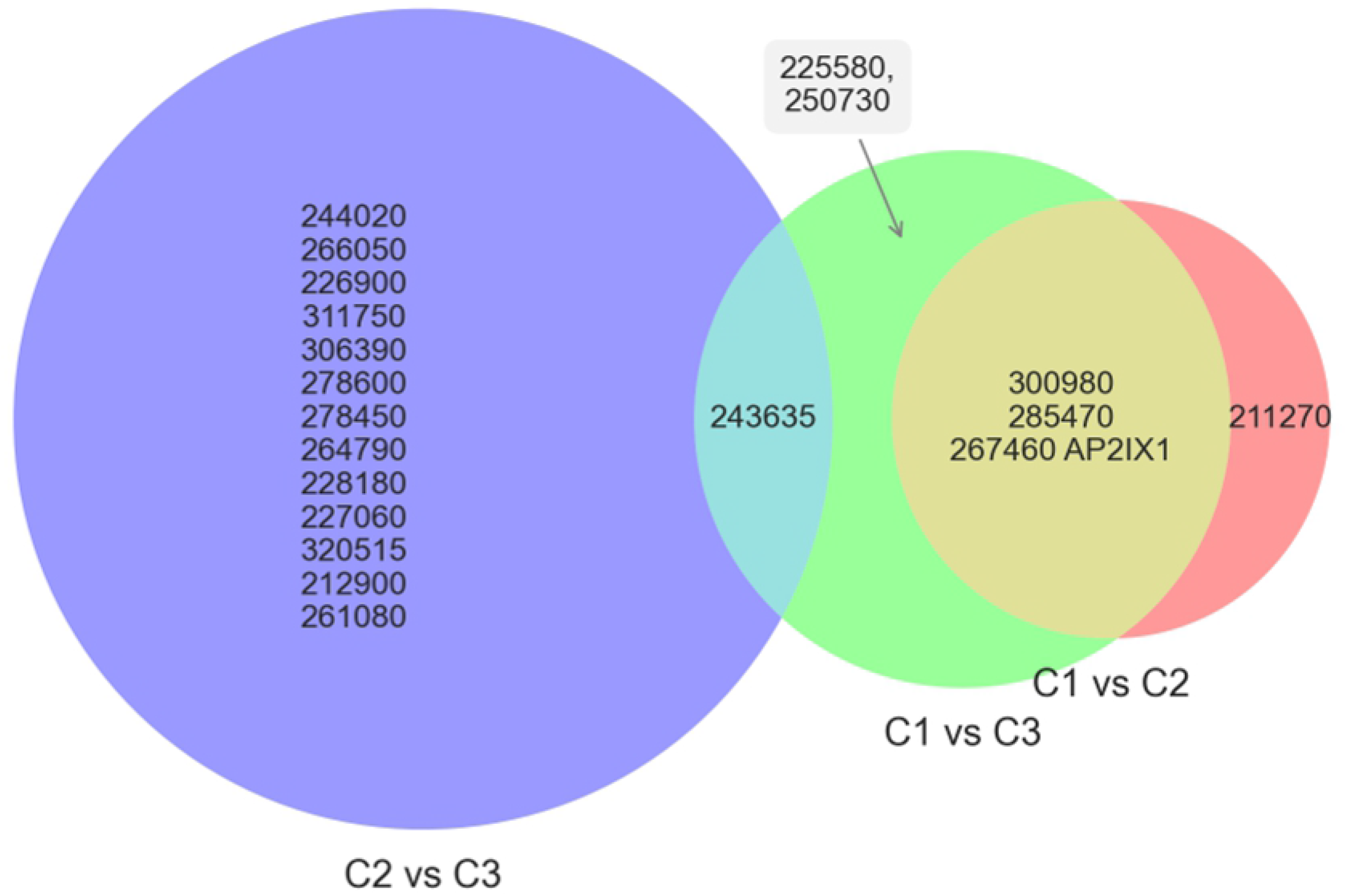
Differentially expressed genes between clonally derived samples. Differentially expressed genes were identified using DESeq2. Pairwise comparisons were made between the different clone lineages by using the subclone data as biological replicates (C1a, C1b, C1c vs C2a, C2b, C2c, etc.). Genes were selected as differentially expressed if their adjusted p-value was less than 0.05 in any of the three pairings (C1 vs C2, C1 vs C3, and C2 vs C3). Genes are shown with their corresponding TgME49 gene ID value. Only one of these genes, 267460, has been annotated with a name (AP2IX1); the rest are of unknown functions or “hypothetical”. No genes were significantly different in all three pairwise comparisons.

To identify any *SRS* expression differences between the subclones (vs. differences between *clones*, discussed above) and to account for the differences in *Toxoplasma* read sums and gene counts between our samples, we randomly chose and down-sampled a constant number of reads from each subclone, based on the lowest read number among the entirety of all samples; i.e., 61,544 reads which was the total for C4. We did this down-sampling five independent times (r1-r5) for each subclone. This reduced the likelihood of the subclones with the highest numbers of reads artifactually appearing more similar to each other due to a higher probability of detecting the same low-abundance genes in them vs. the subclones with lower read depth.

To determine if *SRS* genes were no longer detected in the down-sampled data, which would suggest that *SRS* genes were detected at low read counts and thus their detection in a sample is a product of greater read depth, we compared the amount of *SRS* genes detected with at least one read in the total data to the amount of *SRS* genes with at least one read in the down-sampled data for each of the samples. The C1b and C2b samples had 40 and 39 *SRS* genes detected, respectively, when considering the entirety of the reads. For each of the five down-sampled iterations for these two subclones the number of *SRS* genes detected spanned a range of 30-33 and 29-32, respectively, indicating that down-sampling did indeed result in an ~20-25% decrease in the number of *SRS* genes detected. As expected, when all five down-samplings of a given subclone were pooled, the full repertoire of the 40 or 39 *SRS* genes was restored.

Using these down-sampled datasets, we next examined the relationships between the subclones by performing hierarchical clustering of each down-sampled dataset and plotting the resulting relationships (Fig 5). As expected, the five independently down-sampled replicates for each of the samples were by far the most closely related. The subclones of a given clone, however, were generally no more closely related to each other than they were to unrelated subclones or clones; the C1, C2, and C3 subclones were interspersed, as was the Clone C4.

**Fig 5.**
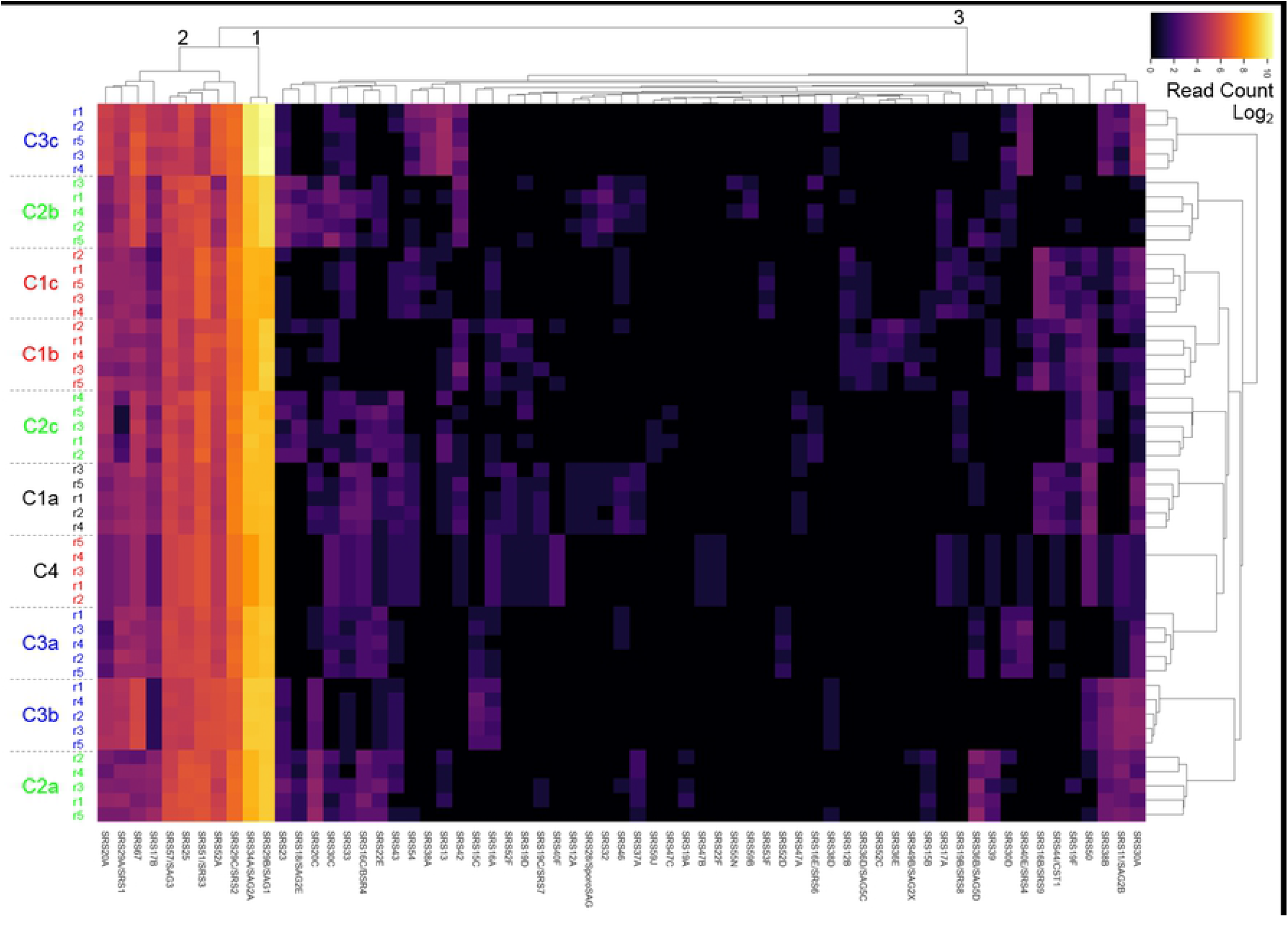
SRS expression among clonally derived tachyzoites. To account for differences in read depth between the samples, the reads for each sample were randomly down-sampled 5 times (r1, r2, etc). The *SRS* genes were selected from these and then the distances between the samples as regards expression levels for each of these *SRS* genes were then calculated. The heatmap shows the relative expression level for each detected *SRS* in all samples, ordered by relatedness between genes (top dendrogram) and samples (right dendogram). The heatmap is log_2_ CPM values for the detected *SRS* genes. Black numbers indicate the groupings of the *SRS* genes.

These data confirm previous observations that *SRS* genes fall into four classes based on their expression data among these tachyzoites samples. In class 1 are the highly expressed *SAG1* and *SAG2A* transcripts, that are among the top 5 most abundantly expressed of all genes and readily detected in all subclones (26). In class 2, there are 9 *SRS* genes that were consistently detected at moderately high expression levels in most samples. Class 3 includes the rest of the *SRS* genes shown in Fig 5 (n=54) that are sporadically detected in at least one but not all of these tachyzoite samples. The final class, class 4, includes the remaining 46 annotated *SRS* genes that were not detected above the required threshold [4 CPM] among any of the samples analyzed here.

## Discussion

Here we present results probing the transcriptional profile of *Toxoplasma* tachyzoites after cloning and subcloning and specifically examine how clonal propagation influences the expression of the *SRS* gene family of surface proteins. Our visual examination and counting at 72 hours after isolating individual parasites indicates between 9 to 11 replicative cycles had occurred resulting in approximately 500 to 2000 parasites. At this timepoint, there did not appear to be a strong and distinct transcriptional pattern that distinguished parasites derived from the same parent clone compared to parasites derived from another parent clone. This indicates that within 10 replications, parasites are expressing transcripts to a similar degree of heterogeneity as seen in parasites generated from a population that has been grown for hundreds of generations. This supports the finding from Xue et al., where nearly all the *SRS* genes were detected across the single-parasite RNA sequencing dataset despite only an average of 7 *SRS* genes being expressed in an individual parasite and suggests that *SRS* gene expression is not stably conferred to progeny over multiple generations (26).

We define here four classes of *SRS* genes based on their expression in these cloned tachyzoite lines: class 1 is ubiquitously and very abundantly expressed (*SAG1* and *SAG2A*); class 2 is consistently but moderately expressed; class 3 is sporadically expressed; and class 4 is unexpressed in all these lines. Our results suggest that there is a somewhat sporadic pattern of expression of class 3 *SRS* genes that results in numerous and different *SRS* transcripts being detected in a population of tachyzoites within relatively few replicative cycles after cloning (<10-12) and that this pattern is not stably inherited upon subcloning. Further studies may reveal a pattern to this apparently random expression of SRS genes that the limited number of clones examined here did not reveal. For example, some developmental control may be operating and “leaky” to different extents in the three clonal lines since the C3 subclones showed a lower number of *SRS* genes being expressed compared to C1 and C2, and they were specifically not expressing two canonical bradyzoite *SRS* genes, *SRS9* and *CST1*. This developmental leakiness was clearly not a major driver of the patterns seen since the various subclones did not generally group together when the totality of the *SRS* gene expression was considered. This conclusion is reinforced by looking at two other canonical *SRS* genes that are developmentally regulated, *SporoSAG* (so-called because it is most highly expressed in sporozoites (42)) and SAG2X (bradyzoite-specific (43)); both showed no obvious pattern with SporoSAG being detectably expressed only in subclones C1a and C2b while SAG2X was on only in C1b. Thus, multiple mechanisms are likely operating in control of these genes’ expression.

Across *Toxoplasma*’s life cycle, over half of the 100+ *SRS* genes are expressed at their highest levels during the sexual stages and the class 3 *SRS* genes include many of these sexual-specific *SRS* genes (24). Thus, the results presented here may indicate that there is some transcriptional leakiness in these tachyzoite populations, perhaps as a result of *in vitro* growth. Validation of our data by examination of parasites derived from animals will be important, though recovering clones and subclones grown for so few generations in animals will be extremely difficult to accomplish. It could be, however, that the results presented here do reflect the *in vivo* reality of an acute infection and that there is a low level, but constant flux, of class 3 *SRS* gene expression in tachyzoites. Indeed, at least in bulk populations, transcripts from many of these class 3 *SRS* genes are detected in tachyzoites from acutely infected mice (44) and this “leaky” expression could have a biological purpose. For example, it could contribute to immune evasion through a mechanism known as “Original Antigenic Sin” that has been well-described for Influenza virus and HIV/AIDS and where related antigens elicit a muted antibody response by presenting closely similar but not identical epitopes (45, 46). Alternatively, it could enable individual tachyzoites to infect a given tissue or host with greater efficiency. Further study of the precise role these proteins have in attachment or other functions in their different hosts and developmental forms will be needed to fully resolve the overall role of the SRS family in *Toxoplasma* biology. The results presented here, however, show that their expression is highly plastic, at least under these conditions.

## Supporting Information

S1 Table: Sample and Read Count Matrix

S2 Table: Sample and log_2_ CPM (Counts per Median) Matrix with gene annotations

S3 Table: DESeq2 Results for differentially expressed genes between samples

## Acknowledgements

The authors especially thank Yuan Xue and Michael Panas for critical review of the manuscript, Melanie Espiritu for help with tissue culturing and ordering, and all members of our, Elizabeth Egan’s, and Ellen Yeh’s labs for helpful comments and suggestions. This work was supported in part by NIH (RO1 AI021423) to JCB and fellowship support to TCT from NIH (T32-AI007328) and Stanford BioX (Stanford Interdisciplinary Graduate Fellowship).

## Author Contributions

Conceptualization: TCT JCB

Data Curation: TCT

Formal Analysis: TCT

Funding Acquisition: TCT JCB

Investigation: TCT

Methodology: TCT JCB

Project Administration: JCB

Resources: TCT JCB

Software: TCT

Supervision: JCB

Validation: TCT

Visualization: TCT JCB

Writing - Original Draft Preparation: TCT JCB

Writing - Review & Editing: TCT JCB

